# Genotyping strategies for detecting CRISPR mutations in polyploid species: a case study-based approach in hexaploid wheat

**DOI:** 10.1101/2021.11.18.469120

**Authors:** Ajay Gupta, Wanlong Li

## Abstract

As a versatile tool for genome engineering, CRISPR-Cas9 has been revolutionizing the field of molecular biology, biotechnology, and crop improvement. By precisely targeting pre-selected genomic sites, CRISPR-Cas9 primarily induces insertions or deletions (indels) of variable size. Despite the significant advance in the technology *per se*, detecting these indels is the major and difficult part of the CRISPR program in polyploid species, like wheat, with relatively low mutation rates. A plethora of methods are available for detecting mutations, but no method is perfect for all mutation types. In this case study, we demonstrated a new, protocol for capturing length polymorphism from small indels using a nested PCR approach. This new method is tractable, efficient, and cost-effective in detecting and genotyping indels >3-bp. We also discussed the major genotyping platforms used in our wheat CRISPR projects, such as mismatch cleavage assay, restriction enzyme assay, ribonucleoprotein assay, and Sanger sequencing, for their advantages and pitfalls in wheat CRISPR mutation detection.

## Introduction

Rapid advance in genome editing technologies has revolutionized the biological sciences and related fields, such as molecular medicine, biotechnology, and crop improvement. Clustered Regularly Interspaced Short Palindromic Repeats (CRISPR) together with CRISPR associated protein 9 (Cas9) has become the most widely used genome editing system due to its tractability. As a two-component system, CRISPR/Cas9 involves a conditional Cas9 nuclease and a single guide RNA (sgRNA) (Doudna et al., 2014). The sgRNA consists of a 76-nucleotide long guide RNA scaffold in its 3’ end for Cas9-binding and a 20-nt long guide RNA in 5’ end for guiding the Cas9/sgRNA complex to search for the target sequence along with the chromosomal DNA (Cong et al., 2013; Deltcheva et al., 2011; Jinek et al., 2012). When an exact match is made between the guide RNA and a target DNA and a Protospacer Adjacent Motif (PAM), primarily 5’-NGG-3’, is present immediately downstream to the DNA target site, Cas9 undergoes a conformational change, which activates endonuclease domains on Cas9. The activated endonuclease domains create a blunt double-strand break (DSB) in the DNA between the third and fourth nucleotide from the 5’end of PAM motif (Jiang et al., 2014; Jinek et al., 2012; Jinek et al., 2014). The DSB is subsequently repaired by the error-prone non-homologous end joining (NHEJ) or by the error-free homologous recombination (HR) (Puchta, 2017; Puchta et al., 1996). While HR leads to gene correction or replacement (Puchta et al., 1996), NHEJ causes insertions, deletions, and other mutations at the cleavage locus (Gao, 2018; Lieber, 2010; Puchta, 2005; Symington et al., 2011).

The predominant use of CRISPR/Cas9 technology in plants is to target genes of interest for knockout via NHEJ for functional analysis and trait improvement (Jaganathan et al., 2018). In common wheat or bread wheat (*Triticum aestivum* L.), the most widely cultivated crop providing ~20% of our daily calorie and proteins, CRISPR/Cas9 has been used to generate novel variations to improve important traits like grain size and yield (Wang et al., 2018; Zhang et al., 2016; Zhang et al., 2019), disease resistance (Wang et al., 2018; Wang et al., 2014; Zhang et al., 2017), pre-harvest sprouting (Abe et al., 2019), low-gluten content (Sánchez◻León et al., 2018), and drought tolerance (Kim et al., 2018). As a hexaploid (2n = 6x =42) species of recent origin with three highly similar (homoeologous) subgenomes A, B, and D (Gornicki et al., 2014), most genes have three homoeologs present on three homoeologous chromosomes and often function redundantly (Appels et al., 2018; Ramírez-González et al., 2018). This requires simultaneous knock-out of the three homeologs to attain the maximum phenotypic effect. (Borrill et al., 2015; Krasileva et al., 2017; Uauy et al., 2017; Wang et al., 2014). Stably integrated CRISPR-Cas9 transgenes transmit to the next generations and stay active. Therefore, multiple mutations in three homeologs can be achieved in subsequent generations (Wang et al., 2018; Zhang et al., 2019). Although CRISPR-Cas9 technology is rapidly advancing, genome modification of wheat using CRISPR-Cas9 still faces certain challenges due to several reasons. First, transforming wheat with the large CRISPR-Cas9 containing cassettes is tedious and inefficient, and yield a small number of T_0_ events (Adamski et al., 2020; Puchta, 2017; Zhang et al., 2019). Second, the mutation frequency at the T_0_ generation is very low as compared to other plants and it is highly unlikely to simultaneously edit all the three homoeologous genes in a single generation (Adamski et al., 2020; Howells et al., 2018; Zhang et al., 2016; Zhang et al., 2019). This requires screening of a large number of plants in the subsequent generations to search desired indels. Third, the polyploid nature of the wheat genome is not only challenging in generating NHEJ mutations but also in detecting and genotyping them. The homoeologous genomes have more than 95% similarity over the coding regions. This offers the advantage that mostly a single sgRNA is sufficient to target three genomes (Arndell et al., 2019) but poses a great difficulty in designing homoeolog-specific primers required for PCR-based mutation detection approaches. Therefore, a cost-effective mutation screening procedure is critical for the timely identification of desirable mutants from the large populations.

Many methods have been developed for detecting the NHEJ mutations in wheat by analysis of the sequences containing the sgRNA-binding site, which are amplified separately for three genomes. While deletions of size >50-bp can be visualized by agarose gel electrophoresis, small deletions are detected by more sophisticated methods such as nuclease assays (including T7 endonuclease 1 (T7E1), ribonucleoprotein (RNP), and restriction enzyme (RE) assays), high-resolution melt curve analysis, Sanger sequencing, and deep sequencing. All these methods offer some advantages and disadvantages over one another. In this article, we are reporting a case study showing the applications, advantages, and disadvantages of different mutation detection platforms. In addition, we propose a novel, two-step nested PCR approach to detect and genotype small indels in hexaploid wheat. Our novel method is easy to operate, reproducible, and cost-effective, which can be applied to other polyploid species for both detecting and genotyping CRISPR induced mutations.

### Nested PCR: a new method to capture the NHEJ mutations

In our previous report (Zhang et al., 2019), we found 95% of the mutations in wheat are due to deletions, 88% of which are larger than 4 bp, and 35% of which are larger than 50 bp. The mutations with a deletion size of 50-bp or greater can be readily detected based on their length polymorphism. These large deletions have better a chance to completely disrupt the gene and therefore are much desired in gene functional studies. Moreover, these mutations are easy to identify as well as genotype compared to smaller indels. We detected more than 15 large deletions in *TaCKX2* and 1 large deletion in each of *TaSPL13* and *TaGW2* showing the length polymorphism by agarose gel electrophoresis.

Deletions smaller than 50 bp account for approximately two-thirds of total NHEJ mutations (Zhang et al., 2019). Compared to the large deletions, detecting the small deletions (<50 bp), particularly those < 10 bp in size, is much more challenging. Polyacrylamide gel electrophoresis (PAGE) can be an option for detecting the small NHEJ mutations because it has been successfully and widely used to detect small indels such as microsatellites (Wang et al., 2003). To avoid the interference of homoeologous polymorphisms in polyploid, homoeolog-specific primers are required. With this, we developed a nested PCR method for screening large populations for the NHEJ mutations in wheat.

Fig. 1a shows the schematic view of the nested PCR approach of mutation detection and genotyping. In this method, the genomic region flanking the sgRNA target is first amplified using homoeolog-specific primers, and the amplicons are analyzed for large deletions using agarose gel electrophoresis. By using 2-3% agarose gel, large deletions of size > 50-bp can be readily detected. For subsequent detection of small mutations, the first-round PCR product is diluted 1:200 with water and used as a template for second-round PCR, which uses primers surrounding the sgRNA target for an amplicon not exceeding 300 bp (Fig. 1a). The second-round amplification does not require the primers to be homoeolog-specific because the template is homoeolog-specific. Thus, a common primer pair can work for the homoeologs from the A, B, and D genomes. Most mutations are heterozygous at detection, and heteroduplexes can be readily detected by PAGE. Thus, a denaturation-renaturation step is included at the end of second-round PCR to enhance the heteroduplex formation. The nested PCR product can be analyzed on 8% PAGE for detecting small indels of size 3 to 50-bp. The heterozygous small indels can be identified very efficiently due to the presence of heteroduplexes in the gel (Fig. 1a). The identification of heteroduplexes makes this method very attractive for detecting novel mutations of small size. Using this approach, we identified 5-bp and 10-bp deletions in *TaSPL13-D* and *TaSPL13-A*, respectively (Fig. 1b and c). The mutations were confirmed using Sanger sequencing (Fig. 1e). We also used this method to genotype the mutant progenies and found that the method is reproducible. The smallest mutation we genotyped using this method till now is a 5-bp deletion, but the resolution power of >8-% polyacrylamide gel can reach 3 bp (Wang et al., 2003).

**Fig. 1.**
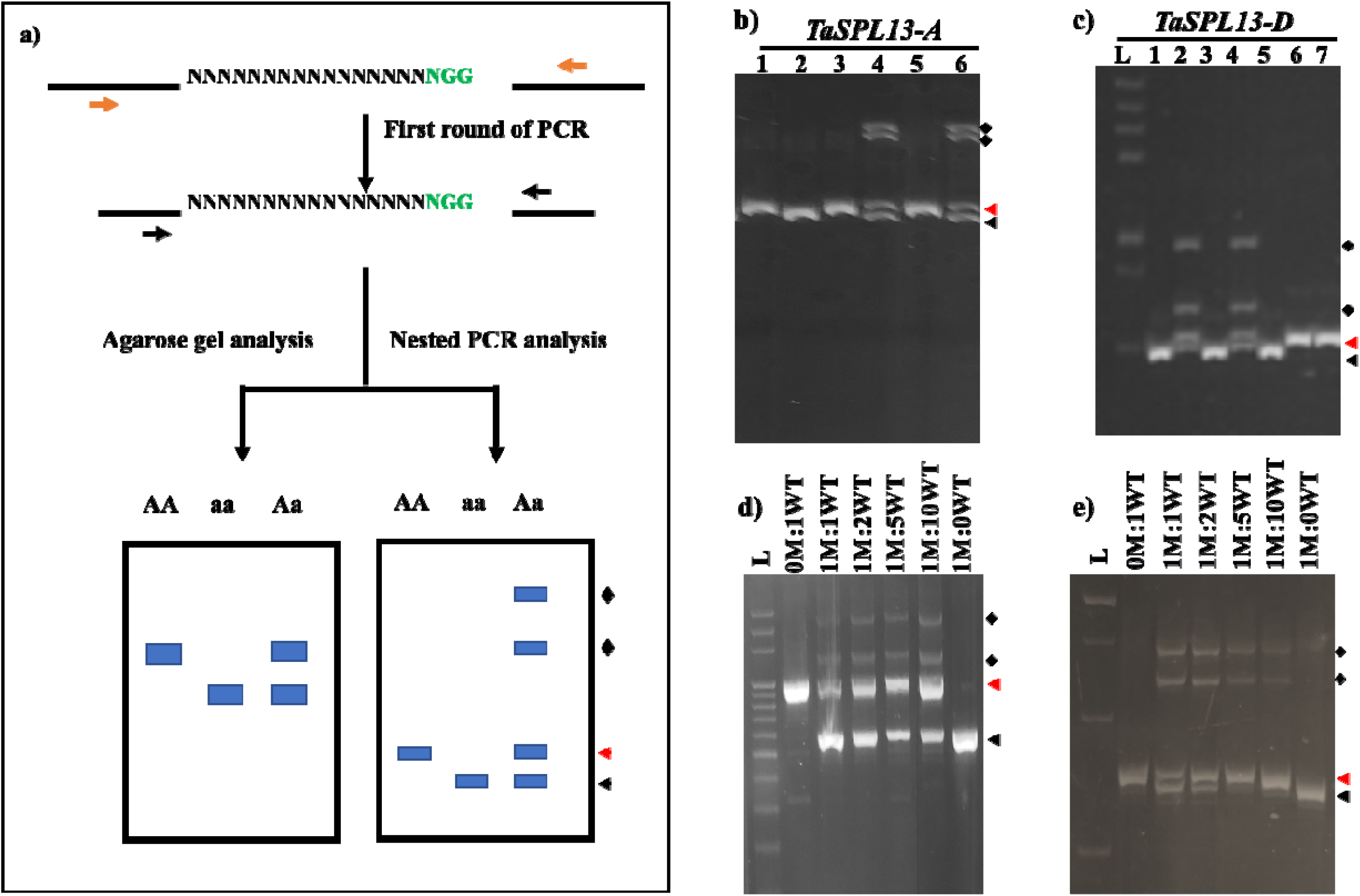
Overall schematic view of the strategy of using a nested PCR approach for detecting large as well as small indels in wheat. (a) Regions targeted by sgRNA (represented by Ns) on three genomes of wheat (A, B, and D) are amplified specifically using genome-specific primers (orange arrows). This PCR product is analyzed on agarose gel for detecting/genotyping large indels. For small indels, the PCR product is analyzed through the nested PCR approach. A common primer for all three gene copies can be used for amplifying a small fragment from the previous PCR product (black arrows). This PCR product is analyzed on PAGE gel for detecting small indels. AA – wildtype allele, Aa – Heterozygous mutant, aa – Homozygous mutant allele. Nested PCR analysis on PAGE gel of b) 10-bp deletion in *TaSPL13-A* and c) 5-bp deletion in *TaSPL13-D*. The upper band in all the pictures is the WT allele and the lower fragment is showing the deletion mutant allele. The curly bracket shows the heteroduplexes formed between the WT and the mutant allele (present only in the heterozygous mutants). d) The nested PCR detected the mutations in mutant:wildtype genomic DNA dilutions of 1:1, 1:2, and 1:5 with 35 cycles of first PCR and 30 cycles of nested PCR amplification. Dilutions are represented on the top of figures. WT = homozygous Wildtype, M = homozygous mutant, L is 100 bp ladder. e) Sanger sequence confirmation of the 10-bp deletion in *TaSPL13-A*, and 5-bp deletion in *TaSPL13-D.* The sequences show the sgRNA target (green), the PAM motif (red), deletion (dashes), and size of mutations (on the right) with the minus sign (−) indicating deletion size.

The choice of two-round PCR over one-round is due to two reasons. Firstly, large deletions can be identified in the first round. Secondly, a common primer pair can be used in the second round because genome-specific primers are difficult to design for a small interval in the coding region due to the very low level of homoeologous variations. The size of the nested PCR product is an important aspect of concern. As the size of PCR products increases, the resolution power decreases, which makes the detection of very small deletions, like 3 to 5-bp deletions, difficult. Thus, for identifying mutations of size less than 8-bp, it is advised to design primers with a product size of <150 bp surrounding the sgRNA, and for an indel of 3-bp a product size of <100 bp is recommended for the nested PCR.

This nested PCR method offers several advantages over the other mutation detection methods. First, it requires no additional enzymes like T7E1, restriction enzymes (RE), or the Cas9 ribonucleoprotein (RNP). Second, it can be universally used for any sgRNA sequence, irrespective of the PCR-RE method, which restricts the selection of gRNA to only certain sequences. Third, a single pair of primers can be designed for simultaneously detecting mutations in the three genomes of wheat. Furthermore, this method is sensitive enough for pooling strategy. The heterozygous mutation can be detected from 2-, 3-, and 5-fold pooled samples (Fig. 1d). Thus, it is a cost-effective, time-saving, and tractable method for detecting NHEJ mutations in polyploid species. To our knowledge, this is a novel method for fast mutation detection in a complex genetic system like wheat.

### Enzyme mismatch cleavage assays

At the early stage of genome editing development, enzyme mismatch cleavage (EMC) analysis was widely used for mutation detection (Cong et al., 2013; Isalan, 2012; Jinek et al., 2013; Woo et al., 2015; Xie et al., 2013; Yang et al., 2017; Zischewski et al., 2017). EMC assay routinely uses T7E1 or Surveyor mismatch cleavage enzymes such as CEL1, CELII, and ENDO1, which have the property to identify, bind, and cleave heteroduplex DNA formed after denaturation and renaturation of wildtype (WT) and mutant alleles (Mashal et al., 1995). Particularly, T7E1 can differentiate between the homoduplex and heteroduplex DNA and specifically target the heteroduplex dsDNA. T7E1 identifies the bulges or kinks in the dsDNA formed due to mispairing of the nucleotides and theoretically can identify a mismatch of even 1-bp (Gohlke et al., 1994). The previous report suggests that the T7E1 is more sensitive and outperforms in mutation detection compared to the Surveyor and thus is the choice of the enzyme for mutation detection (Vouillot et al., 2015).

For EMC assays, a denaturation-renaturation step is included following the completion of PCR amplification. During this denaturation and renaturation step, the PCR products from a heterozygous plant containing both WT and mutant alleles will form the homoduplexes as well as heteroduplex DNA, like nested-PCR of heterozygous mutations (Fig. 2a). In the heteroduplex, the WT and mutant allele DNA anneal throughout their length except at the mutation position, where they mismatch. This mismatched region of the heteroduplex forms a T7E1target site because it produces polymorphic structures including kinks and bulges. The T7E1-digested PCR product can be resolved by agarose gel electrophoresis. For homozygous WT and mutant plants, there will be a single fragment equivalent to the original PCR product in size; for the heterozygous plants, where heteroduplex formation has taken place, three fragments are expected, one large fragment of the original PCR product from undigested homoduplex WT or mutant alleles, and two small fragments due to the digestion of heteroduplex DNA indicative of the presence of the mutation (Fig. 2a). The intensity of digested products is lower compared to the undigested product because the rate of heteroduplex formation is lower than that of homoduplex formation in the mixture (Fig. 2a).

**Fig. 2.**
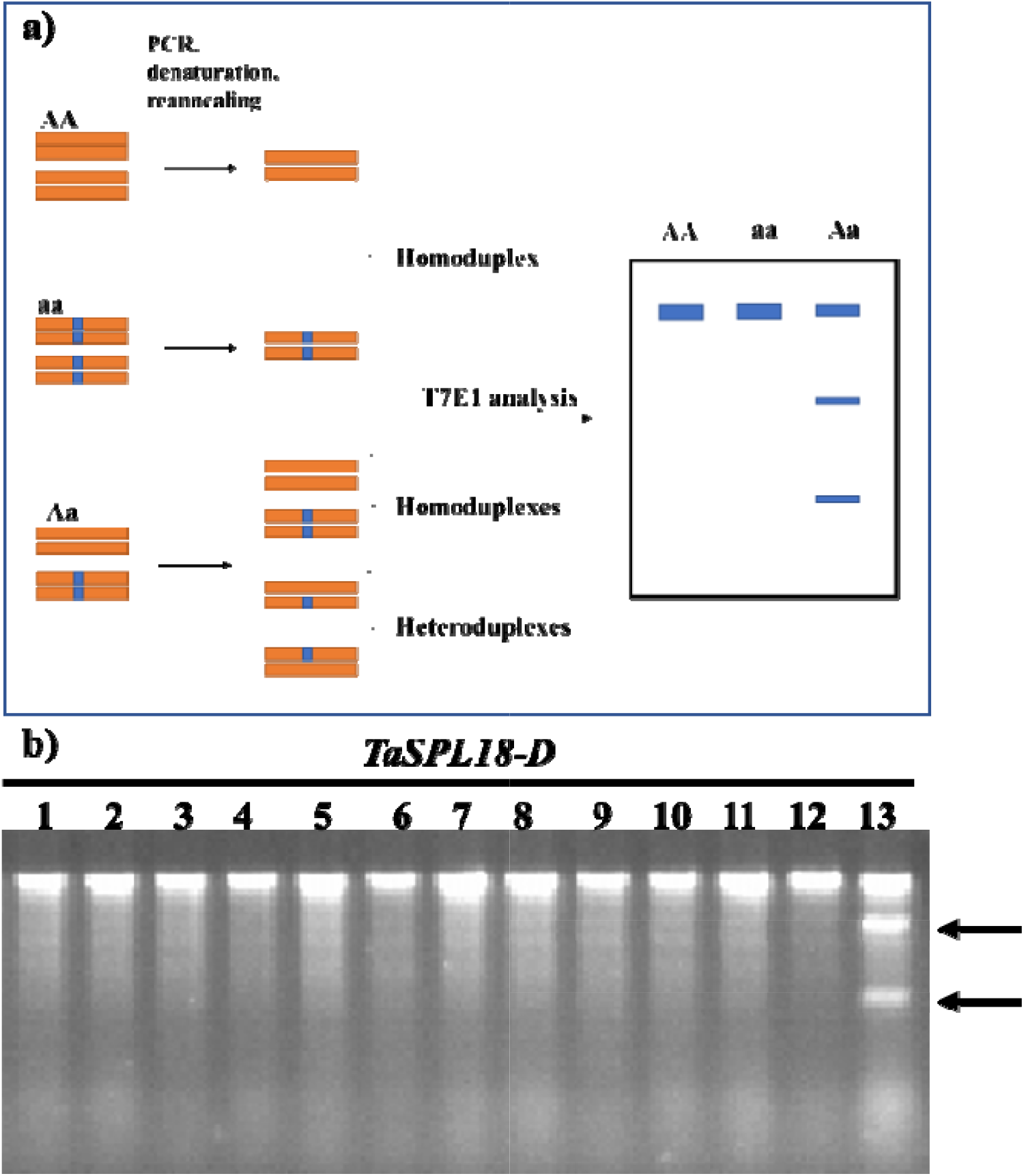
Schematic view of T7E1 enzyme analysis for the detection of mutations. Homozygous WT (AA) and the mutant (aa) alleles remain undigested, while the heterozygous mutant allele (Aa) contains the heteroduplexes that get cleaved into two small fragments (Zischewski et al., 2017). b) A 16-bp deletion in *TaSPL18-D* identified using T7E1 assay. Black arrows indicate the digested bands.

As mentioned previously, T7E1 outperforms other mismatch cleavage enzymes. Thus, we used the T7E1 assay for NHEJ-mutation detections in wheat. The use of T7E1 for mutation detection in wheat requires the target genes to be amplified using genome-specific primers. Primer specificity is very important, otherwise, the inter-genomic variation in the homoeologous genes can be misidentified as NHEJ mutations. We deployed T7E1 for the detection of mutation in two genes targeted by CRISPR/Cas9 including *TaSPL18*, and *TaGW2.* We successfully identified two mutations only in the *TaSPL18* an 83-bp deletion in *TaSPL18-A* and a and 16-bp deletion in *TaSPL18-D* (Fig. 2b). The mutations site and size were confirmed by Sanger sequencing. In the *TaGW2-D* population, the mutation size, 1-bp deletion, was escaped from the T7E1 assay. This is because these 1-bp indels usually occur at the Cas9-cleavage site i.e., between the 3^rd^ and 4^th^ nucleotide from 5’ of PAM and involve an A or T nucleotide (Zhang et al., 2019), but the T7E1 is best in recognizing cytosine nucleotide mismatches (https://www.neb.com/products/m0302-t7-endonuclease-i#Product%20Information).

Despite being extensively used for NHEJ mutation detection, the T7E1 enzyme assay has three major drawbacks. First and most importantly, it is costly, ~$0.5 per reaction. For wheat, where large population screening is required, the method becomes extraordinarily expensive. Secondly, only the heterozygous or biallelic mutations can be identified (Fig. 2a). Thirdly, some small indels, such as 1-bp indels, can be resistant to digestion. Taken together, this method is suitable to screen small populations for detecting novel mutations but not recommended for screening large populations and genotyping of mutations.

### Restriction enzyme assay

RE assay is one of the methods first used to detect CRISPR mutations in wheat (Wang et al., 2014). Because REs can recognize and cleave specific DNA sequences, guide RNA genes must be strategically designed so that the Cas9 cleavage site falls within a RE recognition site. Thus, several guide RNA gene design tools have integrated this function (Cram et al., 2019; Labun et al., 2019) RE assay involves PCR amplification of the sgRNA target site using gene-specific primers, cleavage of the PCR products using the cognate RE, and separation of the digested PCR products by agarose gel electrophoresis. Complete digestion of the WT homozygotes carrying the intact RE site yield two small fragments, but homozygous or biallelic mutants with altered RE site on both the homologous chromosomes are resistant to digestion and yield one undigested, large fragment. The heterozygotes yield three fragments, two small-sized digested fragments from the WT allele and one undigested fragment from the mutant allele (Supplementary fig. S1a). Thus, the PCR-RE assay is a simple and reproducible method well-suited for detecting novel CRISPR mutations and subsequently genotyping. This strategy has been used in several studies to screen the NHEJ mutations in plants (Feng et al., 2013; Shan et al., 2013; Xing et al., 2014). The PCR-RE method is highly sensitive and reproducible. A single-nucleotide polymorphism (SNP) within the RE site is sufficient to abort the RE cleavage site. We utilized this method for detecting several mutations of size ranging from 1-bp to 100-bp in *TaCKX2-3.*1 and *TaGW2-A* (Fig. 3). The sgRNA target site of *TaCKX2-3.1* harbors a restriction enzyme site for *Xcm*I. The RE site of *Xcm*I is 15-nt long and covers a majority of sgRNA targets (Fig. 3a-c and g). We screened 231 T_2_ plants for *TaCKX2-3.1* mutations and identified 30 mutations using *Xcm*I RE. In *TaGW2-A*, a 17-bp deletion aborted the *Xho*I site and we used the *Xho*I site as a codominant marker to genotype the subsequent T_2_ and T_3_ generations (Fig. 3e-g). These results show that the PCR-RE method is highly sensitive and reproducible. In another aspect, PCR-RE assay limits the selection of gRNA because not every target gene contains RE sites. Another factor concerning the use of the PCE-RE method is the cost and availability of REs because routine restriction enzymes are less expensive than rare ones.

**Fig. 3.**
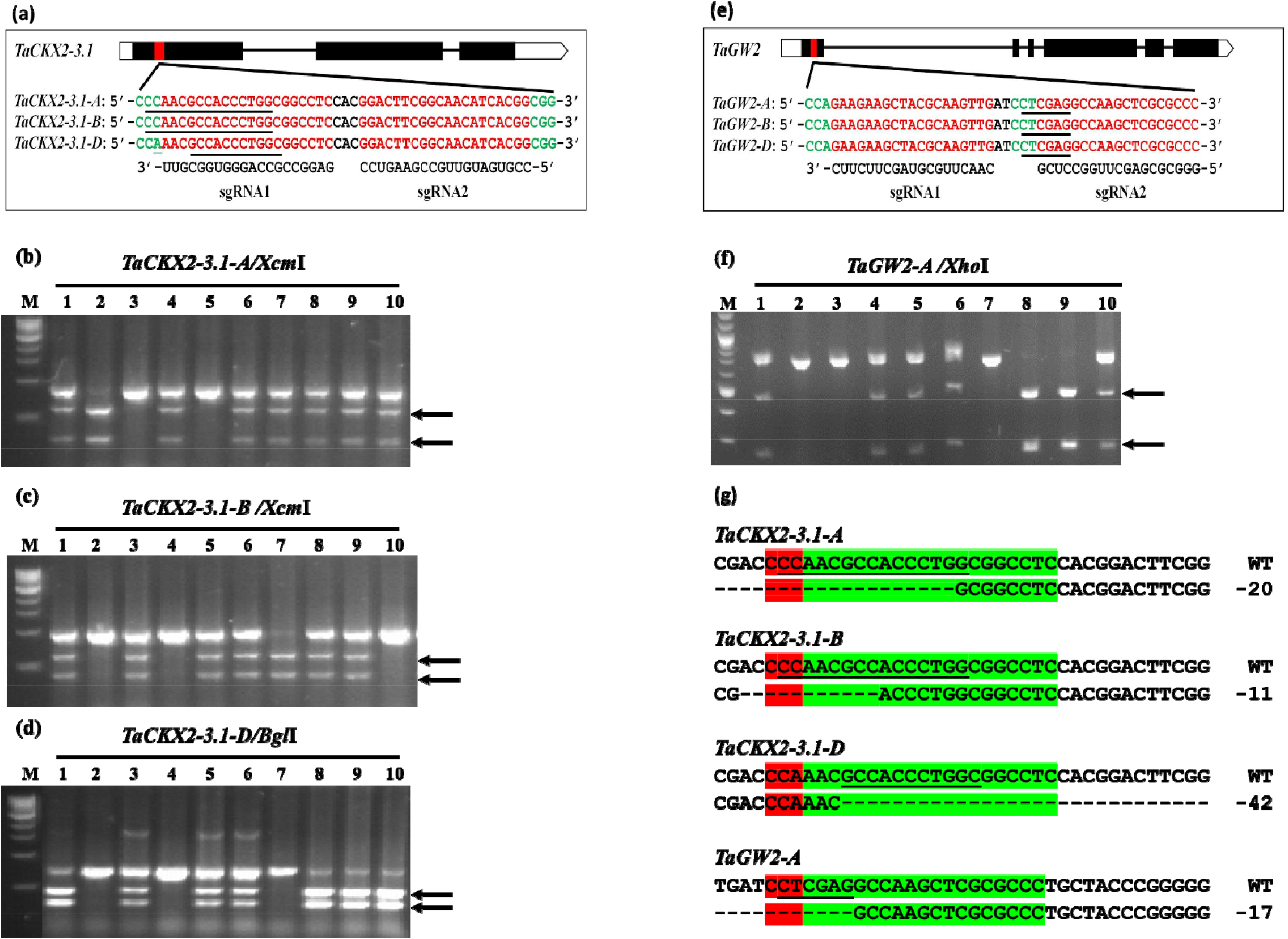
Genotyping of *TaCKX2-3.1* and *TaGW2* using the PCR-RE method. (a) and (e) Gene models of *TaCKX2-3.1* and *TaGW2* and the target sites in exon 1. Red and green nucleotides indicate target sites and PAM sequences, respectively. Underline nucleotides represent the restriction enzyme site (*Xcm*I in *TaCKX2-3.1-A* and *TaCKX2-3.1-B*, *Bgl*I in *TaCKX2-3.1-D*, *Xho*I in all three homoeologous of *TaGW2*. (b) to (d) digestion of *TaCKX2-3.1* homoeologous for mutation genotyping. (f) digestion of *TaGW2-A* with *Xho*I for mutation genotyping. Black arrows indicate at the digested bands. (e) Sequences showing mutations and loss of restriction enzyme site (underlined).

### RNP assay

The creation of DSBs by CRISPR/Cas9 requires an exact match between sgRNA and its target. Once a mutation occurred in the target, CRISPR/Cas9 will no longer recognize the muted target site. Accordingly, the CRISPR/Cas-derived RNA-guided engineered nucleases (RGENs) was initially employed in RFLP analysis by J. M. Kim et al. (2014) and later used for detecting the NHEJ mutations in wheat (Liang et al., 2018). Purified Cas9 protein and *in vitro* transcribed sgRNA are mixed to form the RNP complex, which is used to digest PCR products containing the sgRNA-targeted region. Because the same sgRNA is used for CRISPR and mutation detection, the RNP digests the PCR product of the wild type allele containing an intact sgRNA target site, but it does not cut the mutant PCR product due to disruption of the sgRNA target site by the NHEJ mutations. As a result, one undigested fragment is expected from the homozygous mutants, two small fragments are expected from the homozygous WT plants, and all the three fragments in heterozygous plants contain both the mutant and the wild type allele (Supplementary Fig. S2a and b). Ideally, this method is the most suitable method to detect mutations because it utilizes the sgRNA for mutation detection, and the chance of missing any mutation is theoretically zero.

CRISPR/Cas9 RNP complexes have already been used for DNA free gene editing in human as well as plant cells (S. Kim et al., 2014; Woo et al., 2015). Thus, detailed protocols for purification of Cas9 protein, *in-vitro* transcription of guide-RNA, and PCR-RNP digestion are available (Anders et al., 2014; Liang et al., 2018; Zhang et al., 2019). We utilized the PCR-RNP method to identify novel mutations and for genotyping the mutant populations for *TaSPL13, TaGW2*, and *TaCKX2-3.1* (Fig. 4). We used this method for identifying five novel mutations for *TaCKX2-3.1*, one mutation in *TaGW2-B*, and four mutations in *TaSPL13-B.* All the mutations identified using RNP were confirmed using Sanger sequencing. Also, RNP was successfully used for genotyping these mutations in the subsequent generations (Fig. 4a-f).

**Fig. 4.**
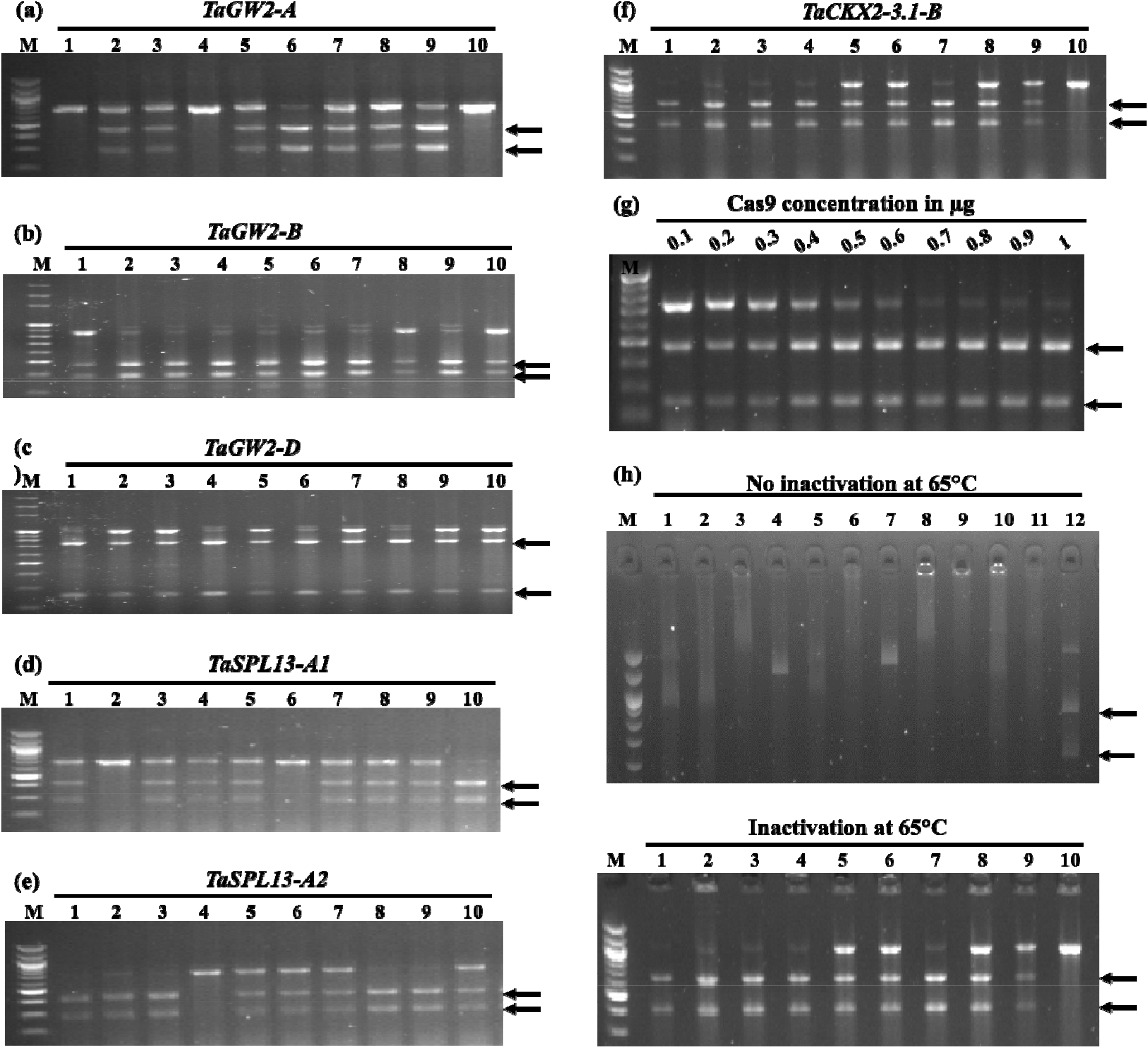
Mutation identification/genotyping of six different genes in hexaploid wheat using PCR-RNP assay and important factors for efficient cleavage. Genotyping of 17-bp deletion in *TaGW2-A* (a), 18-bp deletion in *TaGW2-B* (b), 1-bp deletion in *TaGW2-D* (c), 10-bp deletion in *TaSPL13-A* (d), 25-bp deletion in *TaSPL13-A* (e), and 20-bp deletion in *TaCKX2-3.1-B* (f). Black arrows indicate at the digested bands. (g) Increasing Cas9 concentration from 0.1 to 0.7 μg increases the digestion efficiency, from 0.7 to 1 μg there is no further increase of digestion. (h) Effect of including the 65°C inactivation step after PCR-RNP digestion. No inactivation inhibits the migration of DNA on agarose gel, while digestions inactivated at 65°C migrate normally in the gel.

The cleavage reaction is dependent upon the RNP dosage and the gRNA efficiency. Alone the Cas9 dosage is very important for the complete digestion of the wildtype PCR products (Fig. 4g). The RNP needs to be inactivated at 65°C after the digestion, which separates DNA from the RNP. Otherwise, the DNA-RNP interaction will affect the DNA migration rate (Fig. 4h).

The success of the RNP assay is strongly dependent on the preparation of high-quality Cas9 protein and intact sgRNA. For big population screening, large amounts of Cas9 protein and sgRNAs are required. The commercially available Cas9 proteins are expensive and elevate the cost of genotyping. Expressing the Cas9 in bacteria and purifying manually is an option to minimize the cost of genotyping. The protein isolation and purification protocols are tedious requiring great care to achieve the highest purity and concentration. The sgRNA is also very liable to degradation and requires immense care during handling. Purifying the sgRNA through kits provide better quality and high yield but increases the cost. Low quality and/or low quantity Cas9 and degraded or poor quality sgRNA can lead to false-positive outcomes. The completeness of RNA digestion is determined by the amount of PCR products and the RNA dose used. Thus, overload of PCR products in RNP digestion reactions can yield false-positive results due to incomplete digestion.

### Other methods

In addition to the above-discussed methods, other methods are also, but less commonly, used for detecting mutations, such as deep sequencing, high resolution melting analysis (HRMA), and single-strand conformation polymorphism (SSCP). Deep sequencing is one of the methods of utilizing next-generation sequencing (NGS) technologies to detect mutations. In deep sequencing, the target gene is amplified using PCR and then sequenced using one of the available NGS platforms. An advantage of deep sequencing is that it can detect very low-frequency mutations happening at the somatic cell level. Deep sequencing is a relatively costly method. Pooling the barcoded samples can help reduce the cost of sequencing all samples. Besides, analysis of the massive sequence data requires bioinformatics resources and expertise.

HRMA is a fast, sensitive, and gel-free mutation detection platform (Bassett et al., 2013). This method involves the use of a fluorescent dye which fluoresce brightly when bound to dsDNA and low level when not bound. Thus, upon melting dsDNA the fluorescence brightness is lost and can be detected by a camera. Different dsDNA molecules melt at different temperatures, and in a mixture of PCR amplicons of WT and mutant alleles two peaks of fluorescence can be detected (Thomsen et al., 2012; Wittwer et al., 2003). The 90-150 bp region on the genome targeted by gRNA is amplified using PCR including a fluorescent dye such as SYBR green. The amplified DNA is then melted from 55-95°C while capturing the fluorescence. In the PCR product from the heterozygous plant, two peaks will be indicative of the presence of a mutant allele (Dahlem et al., 2012; Pasay et al., 2008). HRMA is expensive and requires the use of a special qPCR machine to record the fluorescence, thus making the use of this method to resourceful labs.

SSCP utilizes the property of ssDNA to have different conformation due to differences at the sequence level. The ssDNA with different conformation migrates at different rates in PAGE and thus mutations can be identified. The method was first used for detecting DNA polymorphisms between two alleles at chromosomal loci (Orita et al., 1989). This method has been used for mutation detection in rice CRISPR-induced mutants (Zheng et al., 2016). The method can identify genotype small indels of size < 10-bp, but there are two major disadvantages associated with this method. First, the size of the PCR product can’t be more than 250-bp, as the size increases the polymorphism decreases. Secondly, this method has a reproducibility of approximately 80% (Mulhardt, 2010), thus, is not adopted in wheat, in which NHEJ mutation rate is low.

### Sanger sequencing

Except for the NGS, mutations detected by other methods need to be validated by Sanger sequencing. Thus, Sanger sequencing is an indispensable part of CRISPR/Cas9 genome editing and subsequent mutation detection. Most mutations are identified in heterozygous plants. The sequencing chromatograms from heterozygotes contain double peaks starting from the initiation of the indel breakpoint. Resolving this chromatogram to the mutant of unknown sequence and WT sequence manually is time-consuming and error prone. Open access online tools such as DSDecodeM (Liu et al., 2015) (http://skl.scau.edu.cn/dsdecode/) and ICE CRISPR analysis tool (https://ice.synthego.com/) provide platforms to resolve these heterozygous chromatograms to mutant and WT alleles. We deployed Sanger sequencing for validating >70 mutations in *TaCKX2-3.1*, 5 mutations in *TaGW2*, 12 mutations in *TaSPL13*, and 2 mutations in *TaSPL18* genes to determine the size and site of indels (Fig. 5).

**Fig. 5.**
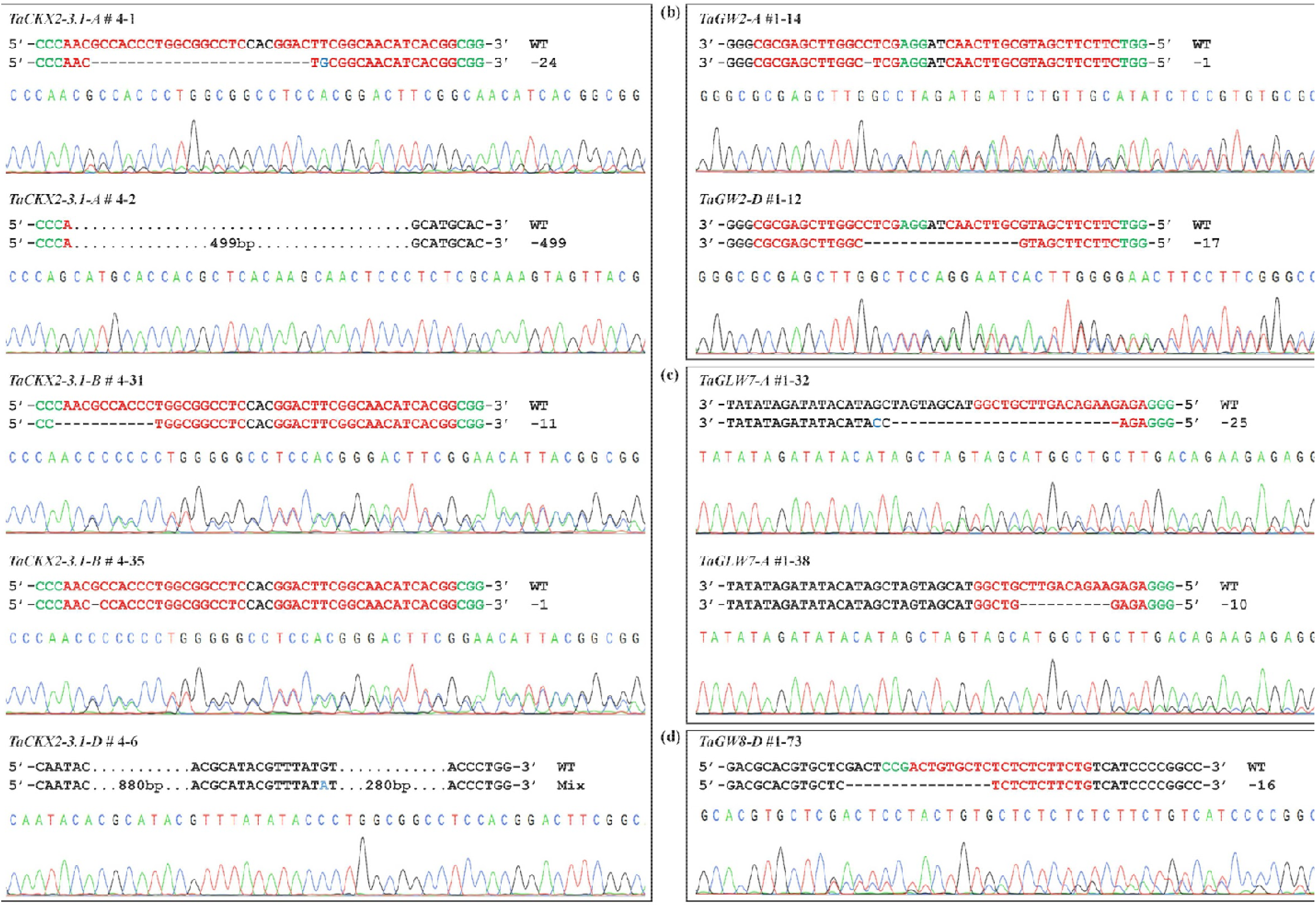
CRISPR/Cas9 induced mutations in target genes. Sequencing chromatograms of PCR amplicons from the selected T_1_ mutants and corresponding regions are shown in (a–d) *TaCKX2*◻*1*, *TaGW2*, *TaSPL13 (TaGLW7)*, and *TaGW8*. The PAM sequences and target sequences are highlighted in green and red, respectively. The dashed lines indicate deletion, and the blue letters represent substitution. Black dots indicate nucleotides not shown (Zhang et al., 2019).

## Conclusion

Except for direct sequencing, a single method is not suitable for capturing the entire spectrum of mutations, and different validation approaches are required to detect novel mutations induced by CRISPR-Cas9. Usually, the number of T_0_ plants obtained after wheat transformation is low. Thus, multiple methods can be utilized to detect novel mutations. Reproducibility and cost-effectiveness are the most important factors to take into consideration while choosing a method for genotyping of known mutations. Therefore, a suitable method, both inexpensive and reproducible, should be selected for the recurrent screening of known mutations in the subsequent generations. Based on our experience, we found the RNP assay, PCR-RE, and nested PCR are the best methods for detecting novel mutations, and nested PCR is the most reliable and inexpensive for detecting and genotyping of mutations of size 3-50 bp.

## Acknowledgments

This research is supported by the USDA Hatch program through the South Dakota Agricultural Experiment Station and USDA NIFA-IWYP (award number: 2017-67008-25934). We are grateful to Dr. Charles Fenster for doing the internal revision of the manuscript.

## Conflict of Interest

The authors declare that they have no conflict of interest.

## Notes

### Competing Interest Statement

The authors have declared no competing interest.

## References cited

Abe, F., Haque, E., Hisano, H., Tanaka, T., Kamiya, Y., Mikami, M., Kawaura, K., Endo, M., Onishi, K., & Hayashi, T. (2019). Genome-edited triple-recessive mutation alters seed dormancy in wheat. Cell reports, 28(5), 1362–1369. e1364.

Adamski, N. M., Borrill, P., Brinton, J., Harrington, S. A., Marchal, C., Bentley, A. R., Bovill, W. D., Cattivelli, L., Cockram, J., & Contreras-Moreira, B. (2020). A roadmap for gene functional characterisation in crops with large genomes: Lessons from polyploid wheat. Elife, 9, e55646.

Anders, C., & Jinek, M. (2014). In vitro enzymology of Cas9. In Methods in enzymology (Vol. 546, pp. 1–20): Elsevier.

Appels, R., Eversole, K., Feuillet, C., Keller, B., Rogers, J., Stein, N., Pozniak, C. J., Choulet, F., Distelfeld, A., & Poland, J. (2018). Shifting the limits in wheat research and breeding using a fully annotated reference genome. Science, 361(6403), eaar7191.

Arndell, T., Sharma, N., Langridge, P., Baumann, U., Watson-Haigh, N. S., & Whitford, R. (2019). gRNA validation for wheat genome editing with the CRISPR-Cas9 system. BMC biotechnology, 19(1), 1–12.

Bassett, A. R., Tibbit, C., Ponting, C. P., & Liu, J.-L. (2013). Highly efficient targeted mutagenesis of Drosophila with the CRISPR/Cas9 system. Cell reports, 4(1), 220–228.

Borrill, P., Adamski, N., & Uauy, C. (2015). Genomics as the key to unlocking the polyploid potential of wheat. New Phytologist, 208(4), 1008–1022.

Cong, L., Ran, F. A., Cox, D., Lin, S., Barretto, R., Habib, N., Hsu, P. D., Wu, X., Jiang, W., & Marraffini, L. A. (2013). Multiplex genome engineering using CRISPR/Cas systems. Science, 339(6121), 819–823.

Cram, D., Kulkarni, M., Buchwaldt, M., Rajagopalan, N., Bhowmik, P., Rozwadowski, K., Parkin, I. A., Sharpe, A. G., & Kagale, S. (2019). WheatCRISPR: a web-based guide RNA design tool for CRISPR/Cas9-mediated genome editing in wheat. BMC Plant Biology, 19(1), 474.

Dahlem, T. J., Hoshijima, K., Jurynec, M. J., Gunther, D., Starker, C. G., Locke, A. S., Weis, A. M., Voytas, D. F., & Grunwald, D. J. (2012). Simple methods for generating and detecting locus-specific mutations induced with TALENs in the zebrafish genome. PLoS genetics, 8(8).

Deltcheva, E., Chylinski, K., Sharma, C. M., Gonzales, K., Chao, Y., Pirzada, Z. A., Eckert, M. R., Vogel, J., & Charpentier, E. (2011). CRISPR RNA maturation by trans-encoded small RNA and host factor RNase III. Nature, 471(7340), 602–607. doi:10.1038/nature09886

Doudna, J. A., & Charpentier, E. (2014). The new frontier of genome engineering with CRISPR-Cas9. Science, 346(6213), 1258096.

Feng, Z., Zhang, B., Ding, W., Liu, X., Yang, D.-L., Wei, P., Cao, F., Zhu, S., Zhang, F., & Mao, Y. (2013). Efficient genome editing in plants using a CRISPR/Cas system. Cell research, 23(10), 1229–1232.

Gao, C. (2018). The future of CRISPR technologies in agriculture. Nat Rev Mol Cell Biol, 19(5), 275–276.

Gohlke, C., Murchie, A., Lilley, D., & Clegg, R. M. (1994). Kinking of DNA and RNA helices by bulged nucleotides observed by fluorescence resonance energy transfer. Proceedings of the National Academy of Sciences, 91(24), 11660–11664.

Gornicki, P., Zhu, H., Wang, J., Challa, G. S., Zhang, Z., Gill, B. S., & Li, W. (2014). The chloroplast view of the evolution of polyploid wheat. New Phytologist, 204(3), 704–714.

Hill, J. T., Demarest, B., Hill, M. J., BiocStyle, S., & biocViews Sequencing, S. (2014). Package ‘sangerseqR’.

Howells, R. M., Craze, M., Bowden, S., & Wallington, E. J. (2018). Efficient generation of stable, heritable gene edits in wheat using CRISPR/Cas9. BMC plant biology, 18(1), 215.

Isalan, M. (2012). Zinc-finger nucleases: how to play two good hands. Nature methods, 9(1), 32–34.

Jaganathan, D., Ramasamy, K., Sellamuthu, G., Jayabalan, S., & Venkataraman, G. (2018). CRISPR for crop improvement: an update review. Frontiers in plant science, 9, 985.

Jiang, W., Brueggeman, A. J., Horken, K. M., Plucinak, T. M., & Weeks, D. P. (2014). Successful transient expression of Cas9 and single guide RNA genes in Chlamydomonas reinhardtii. Eukaryotic cell, 13(11), 1465–1469.

Jinek, M., Chylinski, K., Fonfara, I., Hauer, M., Doudna, J. A., & Charpentier, E. (2012). A programmable dual-RNA–guided DNA endonuclease in adaptive bacterial immunity. science, 337(6096), 816–821.

Jinek, M., East, A., Cheng, A., Lin, S., Ma, E., & Doudna, J. (2013). RNA-programmed genome editing in human cells. elife, 2, e00471.

Jinek, M., Jiang, F., Taylor, D. W., Sternberg, S. H., Kaya, E., Ma, E., Anders, C., Hauer, M., Zhou, K., & Lin, S. (2014). Structures of Cas9 endonucleases reveal RNA-mediated conformational activation. Science, 343(6176), 1247997.

Kim, D., Alptekin, B., & Budak, H. (2018). CRISPR/Cas9 genome editing in wheat. Functional & integrative genomics, 18(1), 31–41.

Kim, J. M., Kim, D., Kim, S., & Kim, J.-S. (2014). Genotyping with CRISPR-Cas-derived RNA-guided endonucleases. Nature Communications, 5(1), 3157. doi:10.1038/ncomms4157

Kim, S., Kim, D., Cho, S. W., Kim, J., & Kim, J.-S. (2014). Highly efficient RNA-guided genome editing in human cells via delivery of purified Cas9 ribonucleoproteins. Genome research, 24(6), 1012–1019.

Kovalchuk, I., Pelczar, P., & Kovalchuk, O. (2004). High frequency of nucleotide misincorporations upon the processing of double-strand breaks. DNA repair, 3(3), 217–223.

Krasileva, K. V., Vasquez-Gross, H. A., Howell, T., Bailey, P., Paraiso, F., Clissold, L., Simmonds, J., Ramirez-Gonzalez, R. H., Wang, X., & Borrill, P. (2017). Uncovering hidden variation in polyploid wheat. Proceedings of the National Academy of Sciences, 114(6), E913–E921.

Labun, K., Montague, T. G., Krause, M., Torres Cleuren, Y. N., Tjeldnes, H., & Valen, E. (2019). CHOPCHOP v3: expanding the CRISPR web toolbox beyond genome editing. Nucleic acids research, 47(W1), W171–W174.

Liang, Z., Chen, K., Yan, Y., Zhang, Y., & Gao, C. (2018). Genotyping genome-edited mutations in plants using CRISPR ribonucleoprotein complexes. Plant Biotechnol J, 16(12), 2053–2062. doi:10.1111/pbi.12938

Lieber, M. R. (2010). The mechanism of double-strand DNA break repair by the nonhomologous DNA end-joining pathway. Annual review of biochemistry, 79, 181–211.

Liu, W., Xie, X., Ma, X., Li, J., Chen, J., & Liu, Y. G. (2015). DSDecode: A Web-Based Tool for Decoding of Sequencing Chromatograms for Genotyping of Targeted Mutations. Mol Plant, 8(9), 1431–1433. doi:10.1016/j.molp.2015.05.009

Mashal, R. D., Koontz, J., & Sklar, J. (1995). Detection of mutations by cleavage of DNA heteroduplexes with bacteriophage resolvases. Nature genetics, 9(2), 177–183.

Mulhardt, C. (2010). Molecular biology and genomics: Elsevier.

Orita, M., Iwahana, H., Kanazawa, H., Hayashi, K., & Sekiya, T. (1989). Detection of polymorphisms of human DNA by gel electrophoresis as single-strand conformation polymorphisms. Proceedings of the National Academy of Sciences, 86(8), 2766–2770.

Pasay, C., Arlian, L., Morgan, M., Vyszenski◻Moher, D., Rose, A., Holt, D., Walton, S., & McCarthy, J. (2008). High◻resolution melt analysis for the detection of a mutation associated with permethrin resistance in a population of scabies mites. Medical and veterinary entomology, 22(1), 82–88.

Puchta, H. (2005). The repair of double-strand breaks in plants: mechanisms and consequences for genome evolution. J Exp Bot, 56(409), 1–14. doi:10.1093/jxb/eri025

Puchta, H. (2017). Applying CRISPR/Cas for genome engineering in plants: the best is yet to come. Current opinion in plant biology, 36, 1–8.

Puchta, H., Dujon, B., & Hohn, B. (1996). Two different but related mechanisms are used in plants for the repair of genomic double-strand breaks by homologous recombination. Proceedings of the National Academy of Sciences, 93(10), 5055–5060.

Ramírez-González, R., Borrill, P., Lang, D., Harrington, S., Brinton, J., Venturini, L., Davey, M., Jacobs, J., Van Ex, F., & Pasha, A. (2018). The transcriptional landscape of polyploid wheat. Science, 361(6403), eaar6089.

Sánchez◻León, S., Gil◻Humanes, J., Ozuna, C. V., Giménez, M. J., Sousa, C., Voytas, D. F., & Barro, F. (2018). Low◻gluten, nontransgenic wheat engineered with CRISPR/Cas9. Plant biotechnology journal, 16(4), 902–910.

Shan, Q., Wang, Y., Li, J., Zhang, Y., Chen, K., Liang, Z., Zhang, K., Liu, J., Xi, J. J., & Qiu, J.-L. (2013). Targeted genome modification of crop plants using a CRISPR-Cas system. Nature biotechnology, 31(8), 686–688.

Symington, L. S., & Gautier, J. (2011). Double-strand break end resection and repair pathway choice. Annual review of genetics, 45, 247–271.

Thomsen, N., Ali, R. G., Ahmed, J. N., & Arkell, R. M. (2012). High resolution melt analysis (HRMA); a viable alternative to agarose gel electrophoresis for mouse genotyping. PloS one, 7(9).

Uauy, C., Wulff, B. B., & Dubcovsky, J. (2017). Combining traditional mutagenesis with new high-throughput sequencing and genome editing to reveal hidden variation in polyploid wheat. Annual Review of Genetics, 51, 435–454.

Vouillot, L., Thélie, A., & Pollet, N. (2015). Comparison of T7E1 and surveyor mismatch cleavage assays to detect mutations triggered by engineered nucleases. G3: Genes, Genomes, Genetics, 5(3), 407–415.

Wang, D., Shi, J., Carlson, S., Cregan, P., Ward, R., & Diers, B. (2003). A low◻cost, high◻throughput polyacrylamide gel electrophoresis system for genotyping with microsatellite DNA markers. Crop science, 43(5), 1828–1832.

Wang, W., Pan, Q., He, F., Akhunova, A., Chao, S., Trick, H., & Akhunov, E. (2018). Transgenerational CRISPR-Cas9 activity facilitates multiplex gene editing in allopolyploid wheat. The CRISPR journal, 1(1), 65–74.

Wang, Y., Cheng, X., Shan, Q., Zhang, Y., Liu, J., Gao, C., & Qiu, J.-L. (2014). Simultaneous editing of three homoeoalleles in hexaploid bread wheat confers heritable resistance to powdery mildew. Nature biotechnology, 32(9), 947.

Wittwer, C. T., Reed, G. H., Gundry, C. N., Vandersteen, J. G., & Pryor, R. J. (2003). High-Resolution Genotyping by Amplicon Melting Analysis Using LCGreen. Clinical Chemistry, 49(6), 853–860. doi:10.1373/49.6.853

Woo, J. W., Kim, J., Kwon, S. I., Corvalán, C., Cho, S. W., Kim, H., Kim, S.-G., Kim, S.-T., Choe, S., & Kim, J.-S. (2015). DNA-free genome editing in plants with preassembled CRISPR-Cas9 ribonucleoproteins. Nature biotechnology, 33(11), 1162–1164.

Xie, K., & Yang, Y. (2013). RNA-guided genome editing in plants using a CRISPR-Cas system. Mol Plant, 6(6), 1975–1983. doi:10.1093/mp/sst119

Xing, H.-L., Dong, L., Wang, Z.-P., Zhang, H.-Y., Han, C.-Y., Liu, B., Wang, X.-C., & Chen, Q.-J. (2014). A CRISPR/Cas9 toolkit for multiplex genome editing in plants. BMC plant biology, 14(1), 327.

Yang, H., Wu, J.-J., Tang, T., Liu, K.-D., & Dai, C. (2017). CRISPR/Cas9-mediated genome editing efficiently creates specific mutations at multiple loci using one sgRNA in Brassica napus. Sci Rep, 7(1), 7489. doi:10.1038/s41598-017-07871-9. (Accession No. 28790350)

Zhang, Y., Bai, Y., Wu, G., Zou, S., Chen, Y., Gao, C., & Tang, D. (2017). Simultaneous modification of three homoeologs of Ta EDR 1 by genome editing enhances powdery mildew resistance in wheat. The Plant Journal, 91(4), 714–724.

Zhang, Y., Liang, Z., Zong, Y., Wang, Y., Liu, J., Chen, K., Qiu, J.-L., & Gao, C. (2016). Efficient and transgene-free genome editing in wheat through transient expression of CRISPR/Cas9 DNA or RNA. Nature communications, 7(1), 1–8.

Zhang, Z., Hua, L., Gupta, A., Tricoli, D., Edwards, K. J., Yang, B., & Li, W. (2019). Development of an Agrobacterium-delivered CRISPR/Cas9 system for wheat genome editing. Plant Biotechnol J, 17(8), 1623–1635. doi:10.1111/pbi.13088

Zheng, X., Yang, S., Zhang, D., Zhong, Z., Tang, X., Deng, K., Zhou, J., Qi, Y., & Zhang, Y. (2016). Effective screen of CRISPR/Cas9-induced mutants in rice by single-strand conformation polymorphism. Plant cell reports, 35(7), 1545–1554.

Zischewski, J., Fischer, R., & Bortesi, L. (2017). Detection of on-target and off-target mutations generated by CRISPR/Cas9 and other sequence-specific nucleases. Biotechnology advances, 35(1), 95–104.

