## Supplemental Fig. S1 and S2 for "Genotyping strategies for detecting CRISPR mutations in polyploid species: a case study-based approach in hexaploid wheat"

### Supplementary file 1

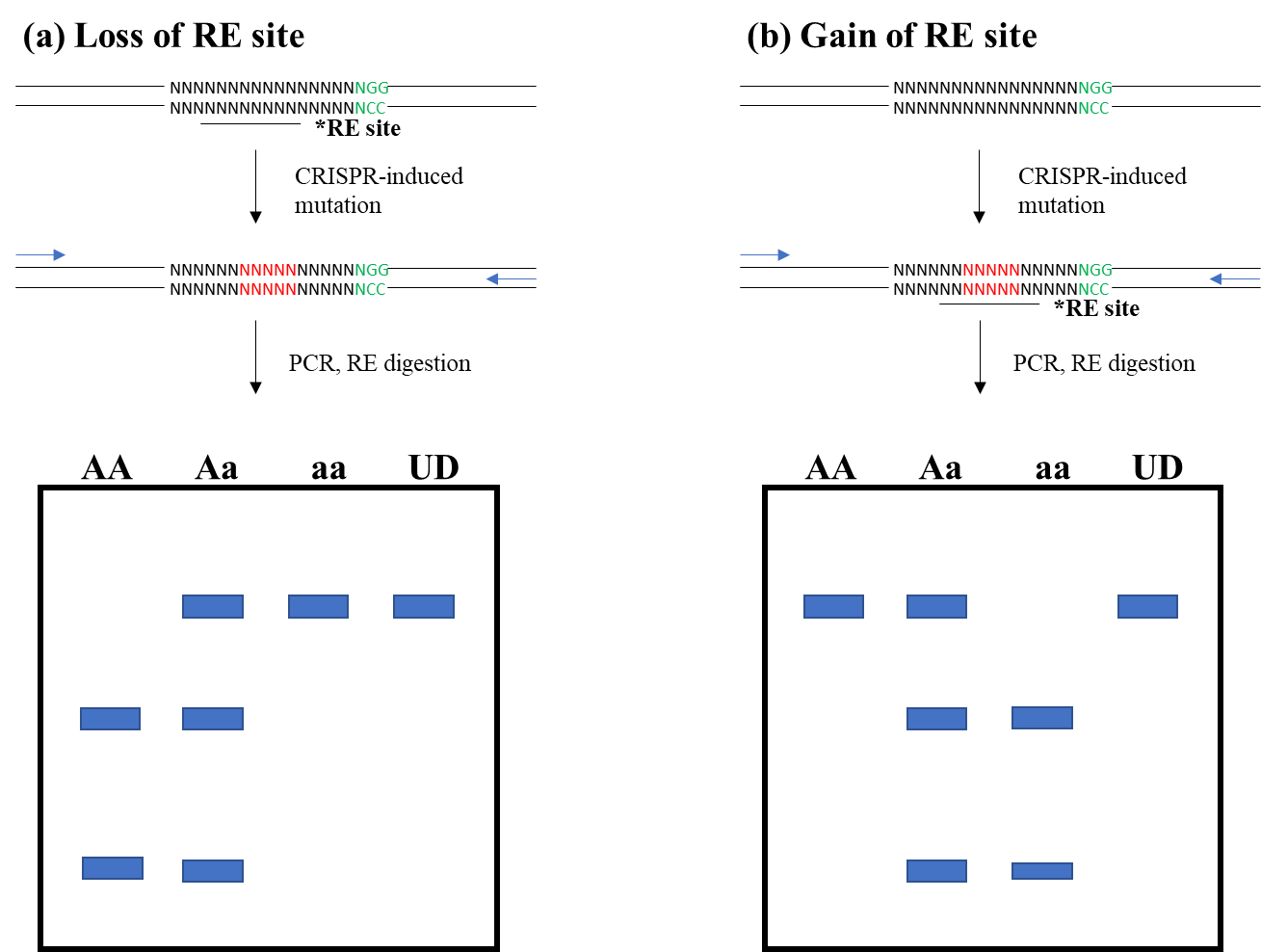

### Fig. S1. Schematic view of using restriction enzyme for detection and genotyping of CRISPR-induced mutations. Guide RNA is represented by Ns followed by NGG motif in green color. Restriction enzyme site is marked by underline (*RE site). (a) Loss of restriction enzyme site upon mutation induction in the sgRNA. WT allele (AA) is digested into two bands, heterozygous mutant allele lane shows three bands, and homozygous mutant allele (aa) remains undigested. (b) The gain of restriction enzyme site upon mutation induction in the sgRNA. WT allele (AA) remains undigested, heterozygous mutant allele lane shows three bands, and homozygous mutant allele (aa) is digested into two bands.

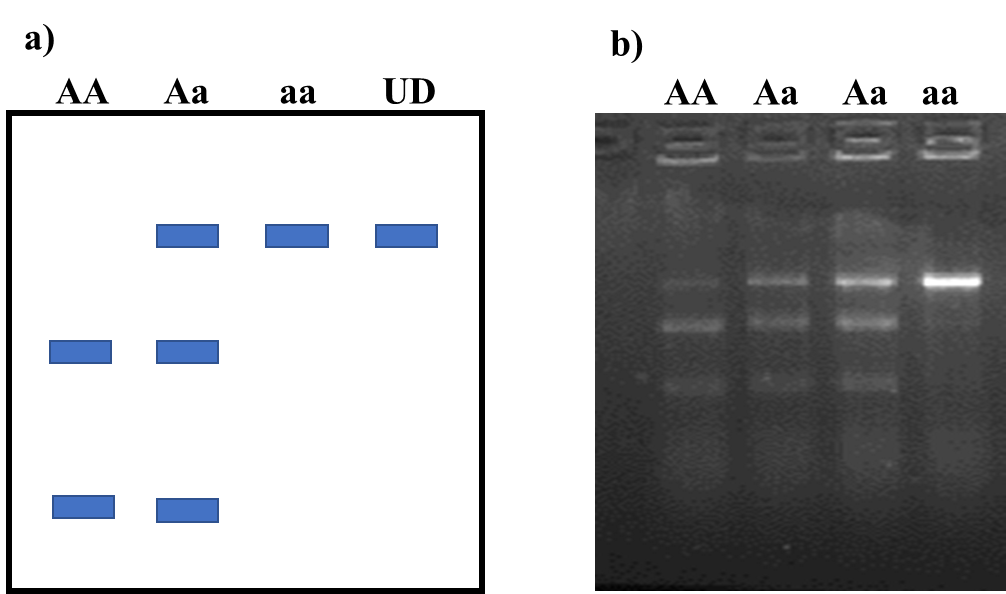

**Fig. S2. Migration of RNP digested PCR product on the agarose gel**. a) Schematic view migration of different outcomes from RNP digestion. b) Digestion of *TaGSK3-A* using the PCR-RNP approach showing different outcomes. AA: Homozygous WT allele, Aa: Heterozygous mutant allele, aa: Homozygous mutant allele, UD: Undigested
